# The suprapyramidal and infrapyramidal blades of the dentate gyrus exhibit different GluN subunit content and dissimilar frequency-dependent synaptic plasticity in vivo

**DOI:** 10.1101/2024.01.08.574587

**Authors:** Christina Strauch, Juliane Böge, Olena Shchyglo, Valentyna Dubovyk, Denise Manahan-Vaughan

## Abstract

The entorhinal cortex sends afferent information to the hippocampus by means of the perforant path(PP), whereby the medial PP (MPP) is believed to convey information about spatial context and the lateral PP (LPP) may convey information about item identity. This information is encoded by means of synaptic plasticity. The PP input to the dentate gyrus(DG) terminates in the suprapyramidal (upper/inner) and infrapyramidal (lower/outer) blades. To what extent frequency-dependent synaptic plasticity in these blades differs is unclear. Here, we compared MPP-DG responses in the supra- (sDG) and infrapyramidal blades (iDG) of freely behaving adult rats and found that synaptic plasticity in the sDG is broadly frequency-dependent whereby long-term depression (LTD, >24h) is induced with stimulation at 1Hz, short-term depression (<2h) is triggered by 5 or 10Hz and long-term potentiation (LTP) of increasing magnitudes is induced by 200 and 400 Hz stimulation, respectively. By contrast, although the iDG expresses STD following 5 or 10Hz stimulation, LTD induced by 1Hz is weaker, LTP is not induced by 200Hz and LTP induced by 400Hz stimulation is significantly smaller in magnitude and is less persistent (<4h) compared to LTP in sDG. Furthermore, the stimulus-response relationship of the iDG is suppressed compared to sDG. Patch clamp recordings, in vitro, revealed reduced firing frequencies in response to high currents, and different action potential thresholds in iDG compared to sDG. Assessment of the expression of GluN subunits revealed significantly lower expression levels of GluN1, GluN2A and GluN2B in iDG compared to sDG. Taken together, these data indicate that synaptic plasticity in the infrapyramidal blade of the dentate gyrus is weaker, less persistent and less responsive to afferent frequencies than synaptic plasticity in sDG. Effects may be mediated by weaker NMDA receptor expression in iDG. These characteristics may explain reported differences in experience-dependent information processing in sDG versus iDG.

## Introduction

The dentate gyrus can be segregated into at least two anatomical subregions: The suprapyramidal (upper/inner) blade (sDG) that is located adjacent to the hippocampal fissure, and the infrapyramidal (lower/outer) blade (iDG) (Amaral and Witter, 1989) (Fig 1A). In vitro examination of granule cell properties suggest heterogeneous membrane and action potential properties are evident within each of the two blades (Mishra and Narayanan, 2020). To what extent information processing at the level of synaptic plasticity differs between the two blades is unclear. However distinctions between the anatomical inputs to sDG and iDG suggest that this might be the case.The primary information source for the dentate gyrus is the perforant path that delivers information from the entorhinal cortex (EC) (Hjorth-Simonsen, 1972; Steward, 1976). The perforant path is segregated into two distinct afferent pathways: the medial perforant path (MPP) that transmits information from the medial EC and synapses at the level of the middle third of the molecular layer, and the lateral perforant path (LPP) that carries information from the lateral EC and synapses at the outer third of the molecular layer (Hjorth-Simonsen, 1972; Steward, 1976; van Groen et al., 2002). It is believed that these pathways provide functionally distinct information to the hippocampus, whereby the MPP path delivers information about the spatial characteristics of sensory experience, whereas the LPP delivers information about no-spatial characteristics such as item identity (Deshmukh and Knierim, 2011; Fyhn et al., 2004; Hargreaves et al., 2005; Kerr et al., 2007; Young et al., 1997).

**Figure 1.**
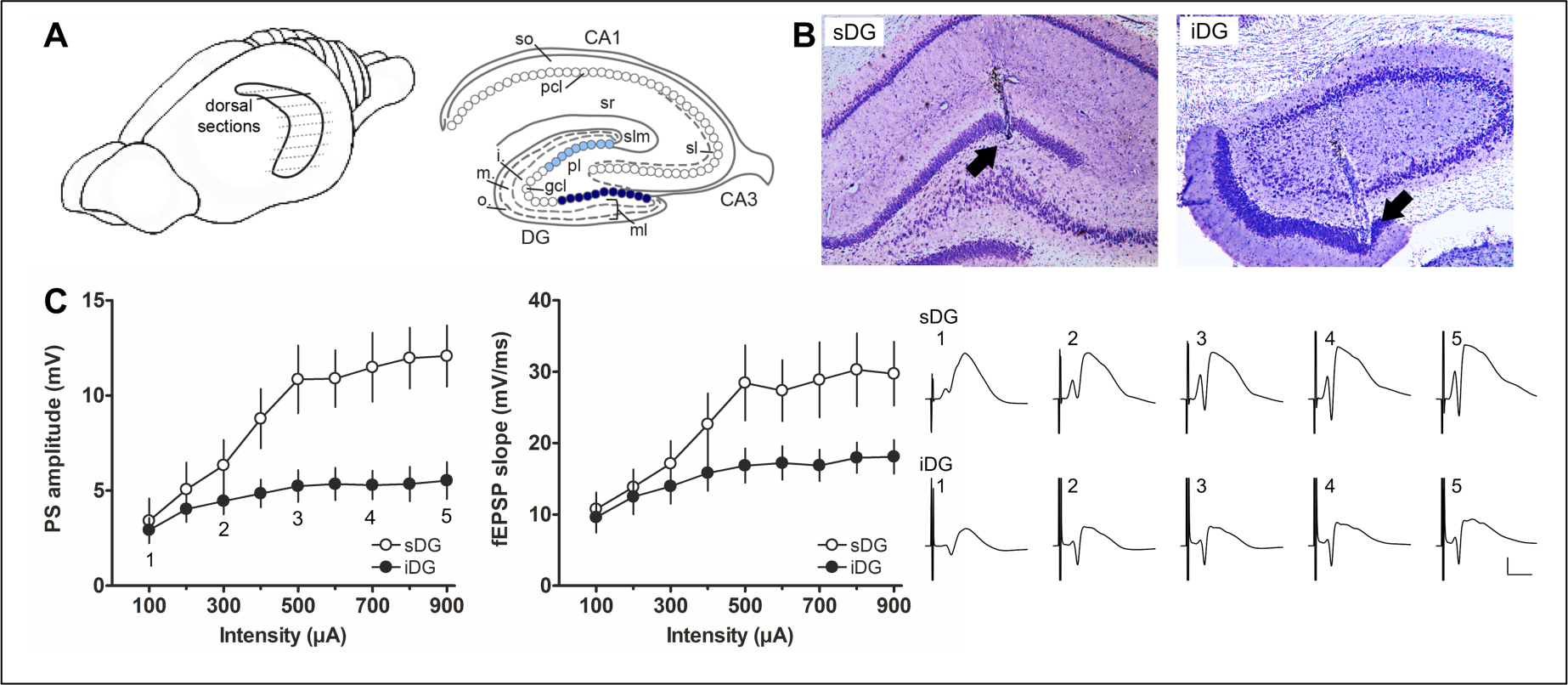
Electrode position and stimulus response relationship of the supra- and infrapyramidal blade of the dentate gyrus. A) Left: Hippocampal formation depicted within a scheme of the rat brain. For immunohistochemistry the brain was sectioned in 30 µm thick horizontal sections (dashed lines) and sections of the dorsal hippocampus (continuous line) were used for analysis. Right: Scheme of the hippocampal formation showing the dentate gyrus (DG) and the cornu ammonis subregions CA1 and CA3 as well as the layers of each subregion. NMDAR subunits were analyzed for the middle molecular layer (m. ml) of the supra- (sDG) and infrapyramidal (iDG) blade of the dentate gyrus. The sDG (light blue) and iDG (dark blue) of the granule cell layer (gcl) are highlighted. B) Left: Nissl-stained coronal section showing the final position of the recording electrode in the granule cell layer of the sDG (ca. 3.1 mm posterior to bregma according to Paxinos and Watson (2014). Right: Example of Nissl-stained coronal section with the final position of the recording electrode in the iDG (ca. 1.9 mm posterior to bregma). The deepest position of each electrode is indicated by an arrow. C) Evaluation of the stimulus response relationship showing PS and fEPSP of simultaneously evoked responses in sDG and iDG (N = 8, each) by test-pulse stimulation of the medial perforant path. Left: PS of evoked responses are significantly different. Middle: fEPSPs show a tendency towards a significance between sDG and iDG. Right: Representative analog examples evoked in each blade at increasing intensities: (1) 100 µA, (2) 300 µA, (3) 500 µA, (4) 700 mA, and (5) 900 µA. Calibration: Vertical bar: 5 mV, horizontal bar: 5 ms.

Anatomical scrutiny has indicated that the lateral EC projects preferentially to the sDG and that the medial EC projects preferentially to the iDG (Tamamaki, 1997; Wyss, 1981). The functional consequence may be that the sDG mainly processes non-spatial, and the iDG predominantly processes spatial information. Proof in support of this prediction is controversial. Although evidence does in fact exist that the sDG and iDG may fulfill different tasks in terms of information processing and storage, scrutiny of experience-dependent expression of immediate early genes after novel exposure to environments of different spatial-/non-spatial contexts suggest that the iDG may be more concerned with the processing of item-place experience and landmark-based directional navigation (Hoang et al., 2018). The sDG, by contrast, may play a role in the processing of spatial navigation in the absence of ostensible landmarks (Chawla et al., 2005), suggesting that both structures contribute to spatial information processing.

Experience-dependent information storage within the hippocampus is mediated by persistent forms of synaptic plasticity (Kemp and Manahan-Vaughan, 2007; Kemp and Manahan-Vaughan, 2008). Although a multitude of studies have scrutinized the frequency-dependency and response characteristics of synaptic plasticity in the dentate gyrus, to our knowledge these studies have focused almost exclusively on the sDG (Altinbilek and Manahan-Vaughan, 2009; Frey and Frey, 2009; Kenney and Manahan-Vaughan, 2013; Krug et al., 1983). To what extent the iDG expresses frequency-dependent synaptic plasticity in vivo has been largely unexplored. Long-term potentiation (LTP) and long-term depression (LTD) have very distinct associations with different kinds of spatial information storage, whereby LTP appears to play an important role in the selection of a neuronal network in the hippocampus (Hoang et al., 2021) and occurs in response to novel spatial experience (Kemp and Manahan-Vaughan, 2004; Kemp and Manahan-Vaughan, 2008), LTD is tightly associated with spatial content learning and content information updating (Goh and Manahan-Vaughan, 2013; Hagena and Manahan-Vaughan, 2011; Kemp and Manahan-Vaughan, 2008; Manahan-Vaughan and Braunewell, 1999). A differentiation in learning-related LTP and LTD in the sDG has been described (Kemp and Manahan-Vaughan, 2008), but whether the frequency-dependency of synaptic plasticity differs between sDG and iDG is as yet unexplored.

The aim of the present study was therefore to compare the frequency-dependency of synaptic plasticity in sDG and iDG in freely behaving adult rats. We chose to analyze plasticity enabled by the MPP as this is the best studied pathway to the dorsal dentate gyrus. Given the importance of the receptor for hippocampal synaptic plasticity (Malenka and Nicoll, 1993; Manahan-Vaughan, 2018; Park et al., 2014), the expression of the GluN1, GluN2A and GluN2B subunits of the N-methyl-D-aspartate receptor (NMDAR) were also scrutinized. Furthermore, using patch clamp recordings in acute brain slices we assessed to what extent granule cell properties differ in the sDG and iDG. We report here that synaptic plasticity in the iDG is weaker, less persistent and less responsive to afferent frequencies than synaptic plasticity in the sDG in vivo. Moreover, action potential properties and firing frequency differs between blades and all three examined GluN subunits exhibit significantly lower expression in the iDG. These differences are likely to potently influence the nature and content of information processing in the iDG.

## Material and Methods

### Subjects

The study was carried out in accordance with the European Communities Council Directive of September 22^nd^, 2010 (2010/63/EU) for care of laboratory animals. All experiments were conducted according to the guidelines of the German Animal Protection Law and were approved in advance by the North Rhine-Westphalia State Authority (Bezirksamt, Arnsberg). The 3R strategy was implemented to reduce the number of animals included in experimentation.

For all experiments male Wistar rats (Charles River, Sulzfeld, Germany) were used. Animals were maintained on a 12h light/12h dark cycle and had *ad libitum* access to water and food. For in vivo experiments, animals were housed individually after surgical implantation of electrodes.

### Surgery

Animals (7-8 weeks old at the time of surgery) underwent chronic implantation of electrodes in the dentate gyrus as described before for the sDG (Manahan-Vaughan and Reymann, 1995). Briefly, electrodes made from polyurethane-coated stainless-steel wire (diameter: 127 µm, Biomedical Instruments, Zöllnitz, Germany) were implanted and screws connected to silver-coated copper wire served as ground and reference electrodes. Anesthesia was implemented using sodium pentobarbital applied intraperitoneally (i.p.) (Nembutal, 52 mg/kg, Boehringer Ingelheim, Ingelheim, Germany). One monopolar recording electrode was placed in the sDG (ca. 3.1 mm posterior to bregma, ca. 1.9 mm lateral from midline, 3.9 to 4.4 mm below the lower edge of the skull) and one in the iDG (ca. 1.9 mm posterior to bregma, ca. 1.0 mm lateral from midline, ca. 5 mm below the skull) according to a stereotaxic rat brain atlas (Paxinos and Watson, 2014). A bipolar stimulation electrode was positioned in the MPP (6.9 mm posterior to bregma, 4.1 mm lateral from midline, 3.8 to 4.2 mm below the skull). After the evoked response in both blades remained stable, the electrode assembly was sealed and fixed to the skull with dental acrylic (Paladur®, Heraeus Kulzer GmbH, Hanau, Germany). Pre- and postsurgical analgesia was implemented using meloxicam in solution (0.2 mg/kg, i.p.; Metacam, Boehringer Ingelheim Vetmedica GmbH, Ingelheim, Germany).

### Electrophysiological recordings

Recording of evoked responses in sDG and iDG were commenced at the earliest ten days after surgery. Evoked responses in the sDG and iDG were evoked by stimulating the MPP at a low frequency (0.025 Hz) using single biphasic square wave pulses (0.2 ms duration per half wave). For all responses, the amplitude of the PS (population spike) and the slope of the fEPSP (field excitatory postsynaptic potentials) were measured.

During all experiments, rats could move freely within the recording chamber (40 x 40 x 50 cm). Disturbances of the animals were kept to an absolute minimum. The stimulus-response (input-output (I/O)) relationship (100 to 900 µA, in 100 µA steps) was recorded to determine the maximum PS. For the remainder of the experiment, a stimulus intensity producing ca. 40% of the maximum I/O response was used to evoke responses. For each time-point five evoked responses were averaged. The first six time-points were recorded at 5 min intervals and these values served as a reference: All data points were calculated as a percentage of the mean of these first six time-points. After recording three further time-points at 5 min intervals, the recording interval was extended to 15 min and a further 15 time-points were recorded. On the next morning, one additional hour of recordings was performed. Rats exhibiting stable evoked responses during a control (test-pulse stimulation) experiment were used for further experiments to assess synaptic plasticity responses. Patterned afferent stimulation (to induce synaptic plasticity) was applied after the recording of the first six reference time-points.

### Patterned afferent stimulation protocols

Patterned afferent stimulation was applied to the MPP at specific frequencies known to induce synaptic plasticity in the sDG (Manahan-Vaughan, 2000; Manahan-Vaughan et al., 1998; Manahan-Vaughan and Reymann, 1995; Twarkowski et al., 2016). Low-frequency stimulation (LFS) at 1 Hz was applied as 900 consecutive pulses. LFS at 10 Hz (450 consecutive pulses) and 5 Hz (10 bursts of 20 pulses with an inter-burst interval of 5 s) were also applied. High-frequency stimulation (HFS) was applied at 200 Hz and 400 Hz. Both protocols consisted of 10 bursts of each 15 pulses applied either at 200 Hz or 400 Hz with an inter-burst interval of 10 s. Recordings of evoked responses were halted during patterned afferent stimulation and recommenced 5 min after the stimulation protocol had concluded. As far as possible, the same animal was used for different plasticity tests. In this case, the sequence of frequency testing was random and at least 7 days elapsed between plasticity experiments. Before animals were re-used, the animal’s I/O responses must have returned to pre-plasticity levels.

### Verification of electrode positions

At the end of the study, brains were removed for histological verification of the position of the electrodes (Fig 1B). Coronal sections (30 µm thick) were stained in 0.1% cresyl violet following a procedure described before (Hansen and Manahan-Vaughan, 2015b). Animals with suboptimally implanted electrodes were excluded from data analysis. In some animals, one of the two recording electrodes was implanted incorrectly, but the data was analyzed for the correctly positioned electrode: therefore, the animal numbers are not equal for all experimental conditions.

### Patch clamp recordings

For acute brain slice preparations, rats (6-8 weeks old) were euthanized by isoflurane anesthesia and decapitation, followed by rapid brain extraction. Horizontal slices (350 µm) were prepared in ice cold, oxygenated sucrose solution (in mM: 87 NaCl, 2.6 MgSO_4_, 75 sucrose, 2.5 KCl, 1.25 NaH_2_PO_4_, 26 NaHCO_3_, 0.5 CaCl_2_, 2 D-Glucose) using a vibratome. Slices were incubated with oxygenated artificial cerebrospinal fluid (aCSF; in mM: 125 NaCl, 3 KCl, 2.5 CaCl_2_, 1.3 MgSO_4_, 1.25 NaH_2_PO_4_, 26 NaHCO_3_, 13 D-Glucose) at 35°C for at least 30 min.

Patch clamp recordings were performed as previously described (Sudkamp et al., 2021). Briefly, slices were continuously perfused with oxygenated aCSF (1-2 ml/min) in a recording chamber under an upright microscope. Borosilicate glass recording pipettes (resistance: 4-7Ma) were filled with an intracellular solution (in mM: 97.5 potassium gluconate, 32.5 KCl, 5 EGTA, 10 Hepes, 1 MgCl_2_, 4 Na_2_ ATP; at pH 7.3). Recordings were conducted from visually identified somata of granule cells of the sDG and iDG. Intrinsic membrane properties were recorded with an amplifier using the PATCHMASTER acquisition software (HEKA Elektronik GmbH, Lambrecht/Pfalz, Germany). After low-pass filtering (2.9 kHz), data were digitized at 10kHz.

After recordings were complete, patched cells were filled with biocytin (1 mg/ml, Sigma-Aldrich, St. Louis, USA) and slices were fixed in 4% paraformaldehyde (PFA) in phosphate buffered saline (PBS) and then kept in 30% sucrose solution for at least 7 days, before being cut into slices of 60 µm thickness at −35°C. Biocytin-filled granule cells were detected using Streptavidin Cy3 (1:1000; Dianova, Hamburg, Germany) (Sudkamp et al., 2021) and nuclei were visualized using 4’,6-diamidino-2-phenylindole (DAPI) in mounting medium (SCR-038448; Dianova, Hamburg, Germany).

### Immunohistochemistry

Immunohistochemistry to assess the expression of the GluN1, GluN2A and GluN2B subunits of the N-methyl-D-aspartate receptor (NMDAR) was performed as previously described (Collitti-Klausnitzer et al., 2021; Dubovyk and Manahan-Vaughan, 2018). Briefly, euthanized animals were transcardially perfused with Ringer’s solution containing heparin (0.2%), followed by 4% PFA in PBS. After removal, brains were fixed in 4% PFA followed by 30% sucrose. Horizontal sections (30 µm thick) were cut, and for each animal, three sections including the hippocampus (ca. 3.6 to 4.1 mm ventral to bregma) were used for immunohistochemistry (Fig 1A). After pretreatment in 0.3% H_2_O_2_, sections were rinsed and incubated with blocking solution containing 10% normal serum (10%), avidin (20%) in PBS with 0.2% Triton X-100 (PBS-Tx). Sections were incubated overnight in primary antibody solution with goat polyclonal anti-NMDAa2 (GluN2B) (1:250; sc-1469, Santa Cruz Biotechnology, Santa Cruz, USA) in 0.2% PBS-Tx with 1% normal serum and 20% biotin. After washing, sections were incubated with biotinylated horse anti-goat (1:500; BA-9500, Vector Laboratories, Burlingame, USA) antibody in 0.1% PBS-Tx with 1% normal serum. Sections were incubated with an avidin-biotin complex (ABC) kit (PK-6100, Vector Laboratories, Burlingame, USA) in 0.1% PBS-Tx with 1% normal serum, after washing. An additional amplification step was performed for other receptors: sections were incubated with primary antibody solutions for 5 days at 4°C: mouse monoclonal antiNMDAR1 (GluN1) (1:200; 556308, PharMingen, Becton, Dickinson and Company, Frankline Lakes, USA) or rabbit polyclonal antiNMDAa1 (GluN2A) (1:750; sc-9056, Santa Cruz Biotechnology, Santa Cruz, USA) in tris-buffered saline containing 0.2% Triton X-100 and 1%BSA. After washing, incubation in biotinylated horse anti-mouse (BA-2001, Vector Laboratories, Burlingame, USA) or goat antirabbit (BA-1000, Vector Laboratories, Burlingame, USA) and further washing, sections were incubated with an ABC reaction. This was followed by the amplification with a biotinylated tyramide solution followed by another ABC reaction. After washing all sections were treated with diaminobenzidine (DAB) and 0.01% H_2_O_2_.

### Data processing and statistics

#### In vivo electrophysiological recordings

For in vivo electrophysiological experiments, data were expressed as a mean percentage of the average reference value and the mean percentage ± SEM of all animals of each group was visualized. To evaluate the I/O relationship, PS and fEPSP values were visualized as mean ± SEM of all animals in which electrodes in sDG and/or iDG were implanted correctly. Using Statistica software (StatSoft. Inc., Tulsa, OK, USA) analysis of variance (ANOVA) with repeated measures was conducted to analyze differences between patterned stimulation and test-pulse stimulation, or to identify differences in effects of the same patterned stimulation protocol on sDG and iDG.

#### Patch clamp recordings

Patch clamp data were analyzed offline using FITMASTER software (HEKA Elektronik GmbH) and AP feature software (MATLAB code provided by Prof. M. Volgushev, Department of Psychology, University of Connecticut). The following action potential (AP) properties were analyzed: AP threshold (mV), spike amplitude from threshold (mV), peak amplitude (mV), afterhyperpolarization (AHP) depth (mV), AHP minimum (mV), total spike time (ms), half-width/width half amplitude (ms), time to peak (ms) and time peak to AHP (ms). In addition, passive and active neuronal properties were assessed: resting membrane potential (mV), input resistance (MOhm), membrane time constant (tau, in ms), excitatory threshold (pA) and firing frequency (Hz). The resting membrane potential was determined from the mean of a 30 s baseline recording. The input resistance was calculated from the slope of the linear fit of the relationship between the change in membrane potential and the intensity of the injected current (−60 pA to 20 pA). Tau was determined from an exponential fit of the averaged voltage decay. The minimum current necessary to evoke an AP from the resting potential was defined as excitatory threshold. Square current pulses (1 s duration) through the patch-clamp electrode (50 pA to 700 pA, 50 pA steps) were applied to examine firing properties. The firing frequency was calculated from the number of spikes elicited during the application of each current step.

Data of patch clamp recordings were visualized using scatter plots including the mean ± SEM of all cells of each blade. The results for the firing frequency were visualized as mean ± SEM of all cells of each blade and were compared using a repeated measures ANOVA. Differences in membrane and AP properties between granule cells of both blades were examined using the Mann-Whitney U test.

#### Immunohistochemistry

Using a light microscope (Leica DMR, Wetzlar, Germany) with a digital camera (MBF Bioscience, Williston, Vermont, USA), photomicrographs of stained sections were acquired and stored in TIFF format to analyze receptor expression. ROIs (regions of interest) of the middle molecular layer of each blade of the dentate gyrus were analyzed at 2.53 lens magnification. Pictures were obtained using Neurolucida software (MBF Bioscience, Williston, Vermont, USA) and quantified using open-source ImageJ software (National Institute of Health, USA). For deconvolution of the color information and conversion to 8-bit format the “Color Deconvolution” plugin in ImageJ was used. Background values from fimbria and deep cerebral white matter tracts were averaged and subtracted from value within the ROI of the corresponding images. For scaling of data from several independent staining/plates, a generalized residual sum of squares algorithm to account for batch effects of staining intensities in R software was used (Kreutz et al., 2007; von der Heyde et al., 2014). These preprocessed values were used for further analysis of immunohistochemistry data: The mean of these values of three sections was calculated for each animal and graphs were created that represent the mean of all animals ± SEM. For statistical analysis, an unpaired t-test was performed to compare the middle compartment of the molecular layer between the two blades for each NMDAR subunit using GraphPad Prism software (version 6, GraphPad Software Inc., USA). The expression patterns of NMDAR subunits of the sDG were published previously (Collitti-Klausnitzer et al., 2021).

For all data, the level of significance was set to p < 0.05. ‘N’ corresponds to the number of animals and ‘n’ describes the number of cells examined in patch clamp recordings.

## Results

### Basal excitability in the infrapyramidal blade is weaker compared to the suprapyramidal blade of the dentate gyrus

To compare synaptic plasticity in sDG and iDG, animals were implanted with one recording electrode in sDG and one in iDG (Fig 1B), to enable examination of the effect of test-pulse stimulation and patterned stimulation on responses evoked simultaneously in both blades of the dentate gyrus.

For comparison of the input-output (I/O) relationship (Fig 1C) only animals (N = 8) were used where both recording electrodes, the one in the sDG and the one in the iDG, were positioned correctly. Here, it was found that the PS of evoked responses in the sDG is significantly different from the iDG (F_(1,14)_ = 6.928, p < 0.05), whereas fEPSPs are not significantly different between the two blades (F_(1,14)_ = 3.149, p = 0.098). However, with increasing stimulus intensities PS and fEPSP of evoked responses in both blades increase (Fig 1C), although this change is weaker in the iDG compared to the sDG (PS: F_(8,112)_ = 11.544, p < 0.0001; fEPSP: F_(8,112)_ = 4.963, p < 0.0001). Post-hoc analysis (Fisher’s LSD test) revealed that the PS becomes significantly different between blades, starting at a stimulus intensity of 400 µA until the maximum intensity that was tested (900 µA), whereas the fEPSP of evoked responses is significantly different at intensities from 500 µA to 900 µA. This difference in the I/O relationship suggests that granule cell excitability differs in the sDG and iDG.

To examine if evoked responses remain stable (Fig 2A), basal synaptic transmission was recorded over a 24h period (4-5 h, followed by 60 min recordings one day after the experiment began). Evoked responses in the sDG (N = 10) and iDG (N = 13) remained stable over time and were not significantly different between the two blades (PS: F_(1,21)_ = 1.765, p = 0.198; fEPSP: F_(1,21)_ = 2.874, p = 0.105).

**Figure 2.**
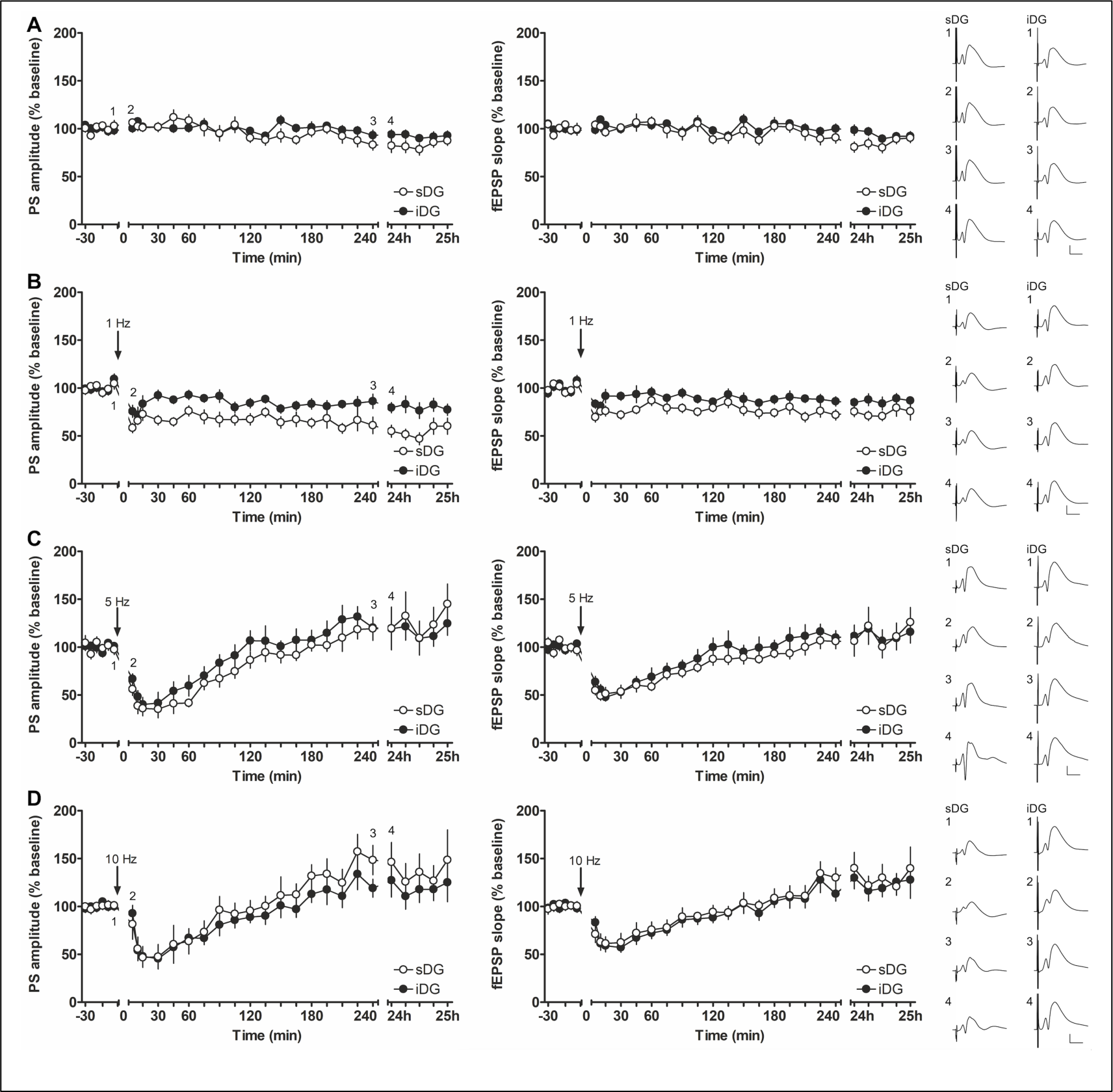
The magnitude of LTD induced by LFS at 1 Hz is weaker in the infrapyramidal compared to the suprapyramidal blade of the dentate gyrus. A) Test-pulse stimulation without application of patterned afferent stimulation evokes stable PS and fEPSPs of field potentials recorded in the suprapyramidal (sDG, N = 10) and infrapyramidal blade (iDG, N = 13) over 4.5h and one hour (24h recording) on the following morning. Analog examples of evoked field potentials recorded in sDG and iDG at the time points (1) −5 min, (2) 5 min, (3) 240 min and during (4) 24h recording. B) Low-frequency stimulation (LFS) at 1 Hz results in (long-term depression) LTD in sDG (N = 9) and iDG (N = 11). LTD induced in the sDG is significantly stronger compared to the iDG. Analog examples of evoked field potentials recorded in each blade at the time points (1) 5 min before, (2) 5 min, (3) 240 min and (4) 24h after LFS at 1 Hz. C) LFS at 5 Hz results in a similar degree of transient depression followed by a potentiation in sDG (N = 8) and iDG (N = 9). Representative examples of evoked field potentials recorded in sDG and iDG at the time points (1) 5 min before, (2) 5 min, (3) 240 min and (4) 24h after LFS at 5 Hz. D) A transient depression followed by a potentiation of a similar magnitude was induced in sDG (N = 7) and iDG (N = 9) by LFS at 10 Hz applied to the medial perforant path. Representative analog examples of evoked field potentials recorded in each blade at the time points (1) 5 min before, (2) 5 min, (3) 240 min and (4) 24h after LFS at 10 Hz. A-D Calibration: Vertical bar: 5 mV, horizontal bar: 5 ms.

### Low-frequency stimulation at 1 Hz induces stronger LTD in the suprapyramidal blade compared to the infrapyramidal blade

Low-frequency stimulation at 1 Hz induces persistent (>24h) LTD in the sDG (Kenney and Manahan-Vaughan, 2013; Manahan-Vaughan, 2000). Here, too LFS at 1 Hz (Fig 2B) triggered LTD in the sDG (PS: F_(1,17)_ = 59.268; p < 0.0001; fEPSP: F_(1,17)_ = 25.362; p < 0.001 versus test-pulse stimulation only). LTD was also induced by 1 Hz (900 pulses) in the iDG (PS: F_(1,22)_ = 16.342; p < 0.001; fEPSP: F_(1,22)_ = 9.829; p < 0.01). A closer comparison of evoked responses of both blades after LFS at 1 Hz (Fig 2B), revealed that LTD in the sDG (N = 9) was stronger than in the iDG (N = 11) (PS: F_(1,18)_ = 17.985; p < 0.001; fEPSP: F_(1,18)_ = 7.234; p < 0.05), suggesting that LTD in the blades of the dorsal dentate gyrus may differ in magnitude in response to 1 Hz stimulation.

### Afferent stimulation at 5 or 10 Hz induces transient depression followed by slow potentiation in the supra- and infrapyramidal blade

Afferent frequencies of 5 Hz or 10 Hz generate transient depression followed by a slowly developing potentiation in MPP-sDG synapses (Twarkowski et al., 2016): Here too, both frequencies evoked a short initial depression followed by a later potentiation in both the sDG (5 Hz: N = 8; 10 Hz: N = 7) and iDG (N = 9, each). Evoked responses after 5 Hz (Fig 2C) and 10 Hz (Fig 2D) were not significantly different between both blades (5 Hz: PS: F_(1,15)_ = 0.585; p = 0.456; fEPSP: F_(1,15)_ = 0.544; p = 0.472; 10 Hz: PS: F_(1,14)_ = 0.589; p = 0.455; fEPSP: F_(1,14)_ = 0.216; p = 0.649). In comparison with baseline experiments conducted using test-pulse stimulation alone, a significant difference was detected for fEPSP values in the sDG after LFS at 5 Hz (F_(1,16)_ = 4.857; p < 0. 05), but not for the PS (PS: F_(1,16)_ = 1.648; p = 0. 218), whereas no significant change can be detected for LFS at 10 Hz (PS: F_(1,15)_ = 1.218; p = 0.287; fEPSP: F_(1,15)_ = 0.820; p = 0.38). For the iDG a comparison with baseline experiments revealed no significant difference for LFS at 5 Hz (PS: F_(1,20)_ = 0.458; p = 0.506; fEPSP: F_(1,20)_ = 2.084; p = 0.164) and at 10 Hz (PS: F_(1,20)_ = 0.325; p = 0.575; fEPSP: F_(1,20)_ = 0.521; p = 0.479). Thus, stimulation of the MPP at 5 or 10 Hz evoked equivalent responses in sDG and iDG.

### LTP induced by HFS at 200 Hz and 400 Hz evoke LTP of greater magnitudes in the suprapyramidal blade compared to the infrapyramidal blade

HFS (Fig 3) applied to the MPP at either 200 Hz, or 400 Hz, evokes persistent (>24h) LTP in the dentate gyrus of freely behaving rats (Manahan-Vaughan et al., 1998; Manahan-Vaughan and Reymann, 1995). We observed that HFS at 200 Hz induces LTP that lasts over 24h in MPP-sDG synapses (N = 9) compared to test-pulse stimulation (PS: F_(1,17)_ = 13.788; p = 0.01; fEPSP: F_(1,17)_ = 21.314; p = 0.001). By contrast 200 Hz HFS elicits no significant change in evoked responses in the iDG (N = 12) in comparison to test-pulse stimulation (PS: F_(1,23)_ = 1.462; p = 0.239; fEPSP: F_(1,23)_ = 3.368; p = 0.079). Evoked responses are also significantly different after 200 Hz HFS (Fig 3A) when both blades are compared (PS: F_(1,19)_ = 10.623; p < 0.01; fEPSP: F_(1,19)_ = 11.796; p < 0.01).

**Figure 3.**
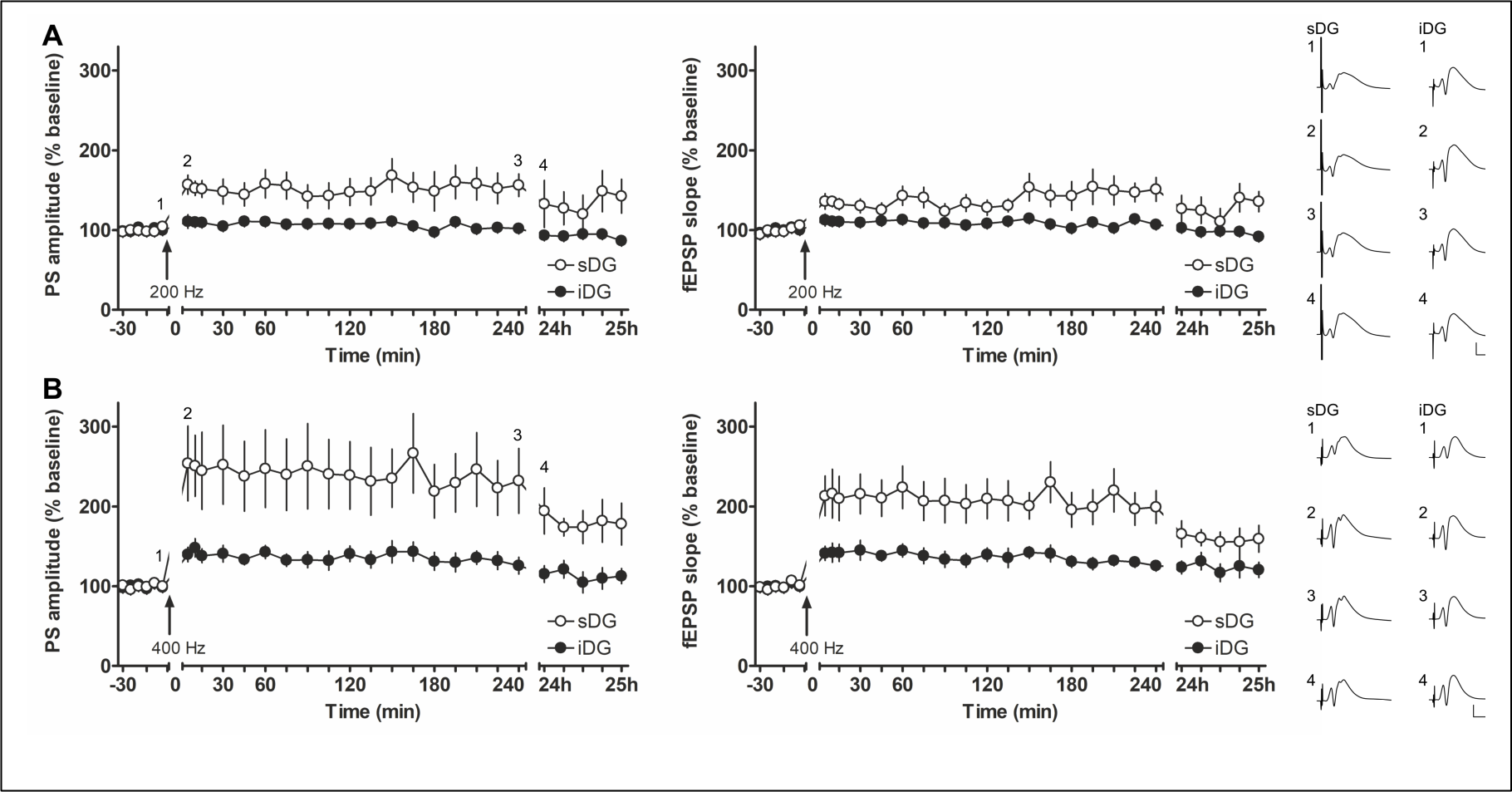
In the infrapyramidal blade, HFS at 200 Hz is ineffective in changing synaptic transmission and 400 Hz HFS induces LTP of a smaller magnitude compared to LTP induced in suprapyramidal blade evoked responses. A) HFS at 200 Hz results in LTP in the suprapyramidal blade (sDG, N = 9) that is significant from the unchanged synaptic transmission induced in the infrapyramidal blade (iDG, N = 12). Analog examples of evoked field potentials recorded in the sDG and iDG at the time points (1) 5 min before, (2) 5 min, (3) 240 min and (4) 24h after HFS at 200 Hz. B) LTP induced by HFS at 400 Hz of the medial perforant path induces a weaker LTP in the iDG (N = 11) compared to the significantly stronger LTP induced in the sDG (N = 9). Representative examples of evoked field potentials recorded in the sDG and iDG at the time points (1) 5 min before, (2) 5 min, (3) 240 min and (4) 24h after HFS at 400 Hz. A-B Calibration: Vertical bar: 5 mV, horizontal bar: 5 ms.

Increasing the afferent frequency to 400 Hz resulted in LTP in iDG synapses (N = 11) compared to test-pulse stimulation (PS: F_(1,22)_ = 18.922; p = 0.001; fEPSP: F_(1,22)_ = 22.932; p < 0.0001). This LTP was smaller in magnitude than responses induced by 400Hz HFS in the sDG (N = 9) in comparison to test-pulse stimulation (PS: F_(1,17)_ = 14.372; p < 0.01; fEPSP: F_(1,17)_ = 31.559; p < 0.0001). Comparison of the LTP responses in both blades (Fig 3B) confirmed that LTP in iDG was significantly smaller than LTP in sDG (PS: F_(1,18)_ = 7.827; p < 0.05; fEPSP: F_(1,18)_ = 11.3; p < 0.01).

Overall, these results suggest that synaptic depression as well as synaptic potentiation evoked by patterned afferent stimulation of the perforant path are substantially weaker in the iDG. Which leads to the questions as to whether these differences arise from blade-specific granule cell properties and/or differences in the expression of plasticity-related receptors in the two blades.

### Distinct action potential properties differ between the supra- and infrapyramidal blades

Closer scrutiny of granule cell properties in the slice preparation revealed no difference between blades for all basic membrane properties (Table 1, Supplementary Fig 1A-D; N = 6 each, n = 19 sDG, n = 22 iDG). However, the firing frequency (Fig 4D-E) differed significantly between blades (100 to 700 pA: F_(1,39)_ = 17.108; p < 0.001). Starting from 350 pA, granule cells in the iDG exhibited a significantly smaller firing frequency compared to granule cells of the sDG (post-hoc: Fisher’s LSD test) demonstrating that the maximal frequency to fire action potentials differs between granule cells in both blades.

**Figure 4.**
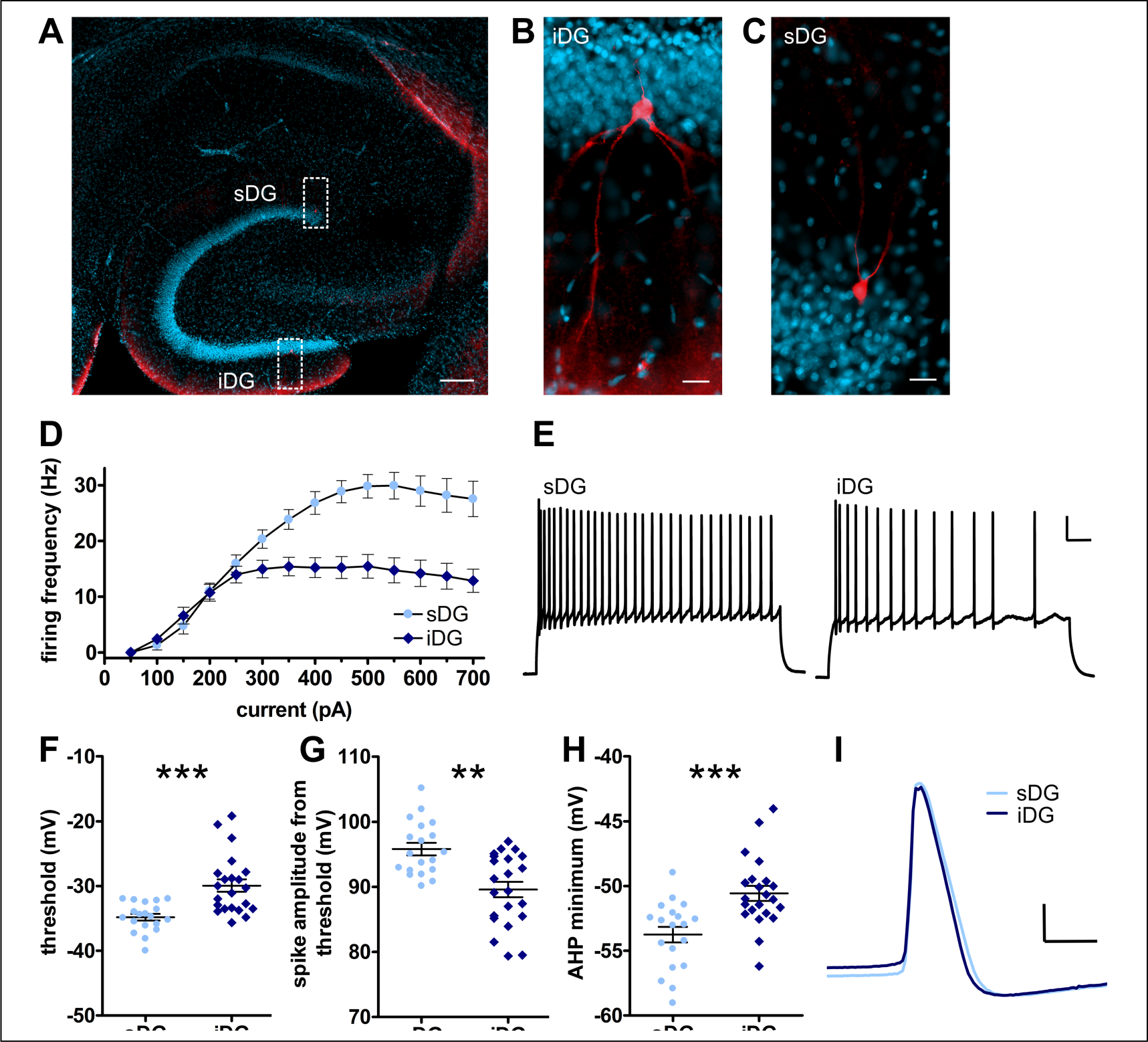
In vitro examination of the firing frequency and action potential properties exhibits differences between granule cells of the supra- and infrapyramidal blade of the dentate gyrus. A-C) Biocytin filled granule cells in the blades of the dentate gyrus. A) Example originating from patch clamp recordings representing the hippocampus in a horizontal section. Granule cells in the sDG and iDG were examined and filled with biocytin (red). Nuclei are counterstained with DAPI (blue). Calibration: 200 µm. B-C) Enlarged images of example in (A) of biocytin filled granule cells in B) iDG and C) sDG. Calibration: 20 µm. D-E) The firing frequency differs between of granule cells of the infrapyramidal (iDG) and suprapyramidal (sDG) blade. D) In granule cells of the iDG (n = 22, N = 6) the firing frequency is significantly smaller compared to granule cells recorded in the sDG (n = 19, N = 6). E) Exemplary traces of the firing frequency recorded at 400 pA in a granule cell of the sDG and one in the iDG. Calibration: Vertical bar: 20 mV, horizontal bar: 100 ms. F-I) Specific action potential properties differ in granules cells of the sDG (n = 19, N = 6) and iDG (n = 22, N = 6). F) The threshold to induce an action potential is significantly higher in the iDG compared to the sDG. G) The amplitude of the action potential from the threshold is significantly smaller in the iDG compared to the sDG. H) The minimum of the afterhyperpolarization (AHP) is significantly lower in the sDG compared to the iDG. I) Example of an action potential recorded in granule cells of the sDG (pale blue) and iDG (dark blue). Calibration: Vertical bar: 20 mV, horizontal bar: 1 ms. F-H) Significant differences are marked with asterisks: ** p < 0.01; *** p < 0.001 Basic membrane properties and further action potential properties are shown in supplementary figure 1.

**Table 1.**
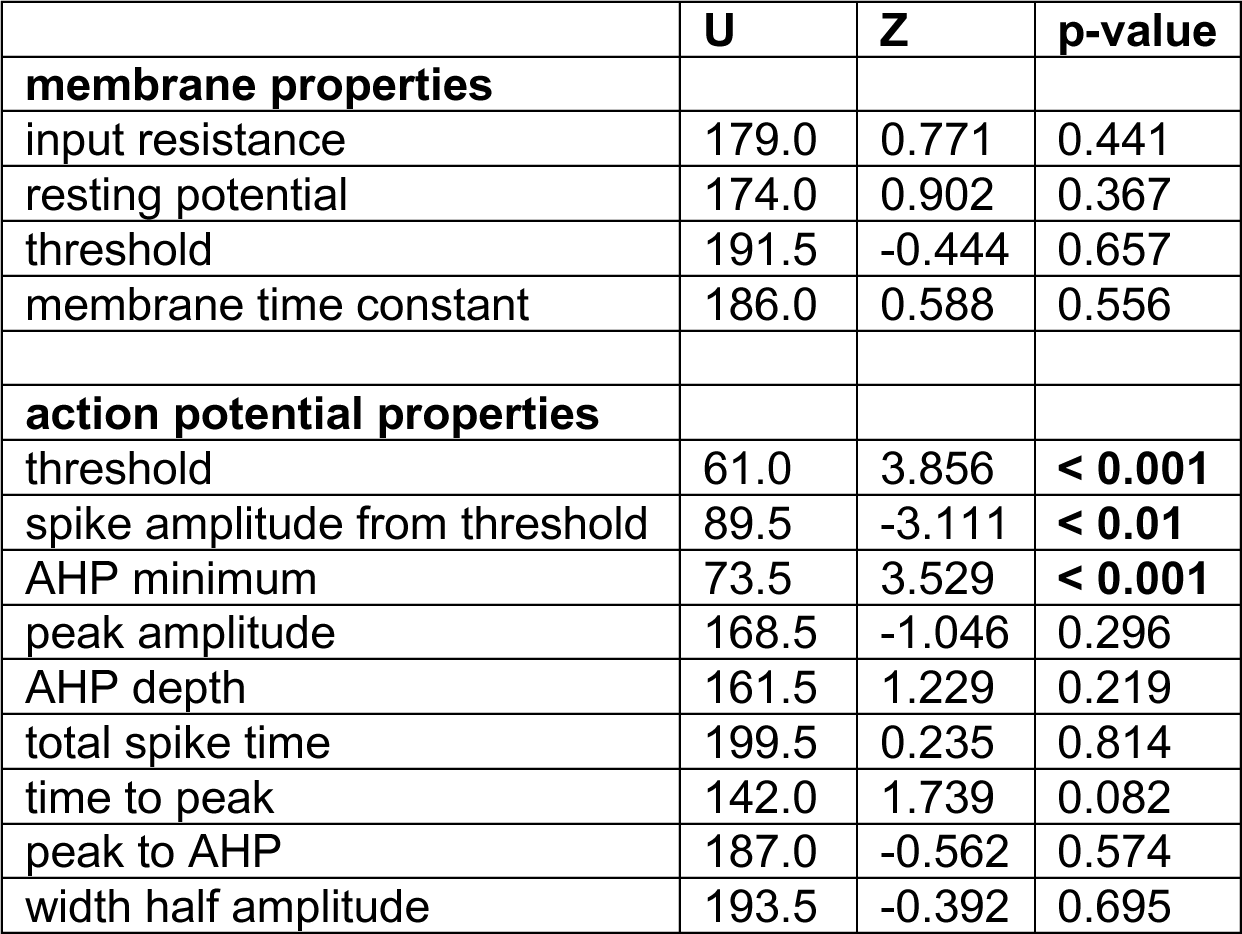
Summary of statistical analysis of basic membrane and action potential properties. Mann-Whitney U test was performed to examine differences in membrane and action potential properties in patch clamp recordings of acute brain slices between infra- (iDG) and suprapyramidal (sDG) blade of the dentate gyrus (N = 6 each, n = 22 for iDG, n =19 for sDG).

|  | U | Z | p-value |
| --- | --- | --- | --- |
| **membrane properties** |  |  |  |
| input resistance | 179.0 | 0.771 | 0.441 |
| resting potential | 174.0 | 0.902 | 0.367 |
| threshold | 191.5 | -0.444 | 0.657 |
| membrane time constant | 186.0 | 0.588 | 0.556 |
| **action potential properties** |  |  |  |
| threshold | 61.0 | 3.856 | < 0.001 |
| spike amplitude from threshold | 89.5 | -3.111 | < 0.01 |
| AHP minimum | 73.5 | 3.529 | < 0.001 |
| peak amplitude | 168.5 | -1.046 | 0.296 |
| AHP depth | 161.5 | 1.229 | 0.219 |
| total spike time | 199.5 | 0.235 | 0.814 |
| time to peak | 142.0 | 1.739 | 0.082 |
| peak to AHP | 187.0 | -0.562 | 0.574 |
| width half amplitude | 193.5 | -0.392 | 0.695 |

Moreover, examination of action potential properties revealed differences between granule cells in the two blades (Fig 4F-I, Table 1): Thus, the threshold to induce an action potential (Fig 4F) was significantly higher in the iDG compared to the sDG (U = 61.0; Z = 3.856; p < 0.001), whereas the peak AP amplitude (Supplementary Fig 1E) did not differ between blades. However, the AP spike amplitude calculated from its threshold (Fig 4G) was significantly larger in the sDG compared to the iDG (U = 89.5; Z = −3.111; p < 0.01). Examination of the afterhyperpolarization (AHP), revealed that the minimum (Fig 4H) was significantly lower in the sDG compared to the iDG (U = 73.5; Z = 3.529; p < 0.001). By contrast the AHP depth is similar in both blades (Supplementary Fig 1F; U = 161.5. All other examined action potential properties, such as the total spike time, or the width half amplitude, were not significantly different between blades (Table 1, Supplementary Fig 1G-J).

### Subunit contents of GluN1, GluN2A and GluN2B receptors are significantly lower in the infrapyramidal compared to the suprapyramidal blade of the dorsal dentate gyrus

We then examined the relative content of subunits of the NMDAR, namely GluN1, GluN2A and GluN2B (Fig 5) in the middle molecular layer (Fig 1A), the region where the MPP synapses (van Groen et al., 2002). Examination of GluN1 (Fig 5 A) revealed a higher expression in the middle molecular layer of the sDG compared to the iDG (unpaired t-test: t_(36)_ = 3.340; p < 0.01; N = 19 each). A similar result was detected for GluN2A (Fig 5B; N = 20 each) and GluN2B (Fig 5C; N = 20 each) subunits (unpaired t-tests: GluN2A: t_(38)_ = 2.745; p < 0.01; GluN2B: t_(38)_ = 5.962; p < 0.0001). Expression of all three subunits was higher in the middle molecular layer of the sDG compared to the iDG. This expression profile may explain why LTP was more potently expressed in sDG compared to iDG.

**Figure 5.**
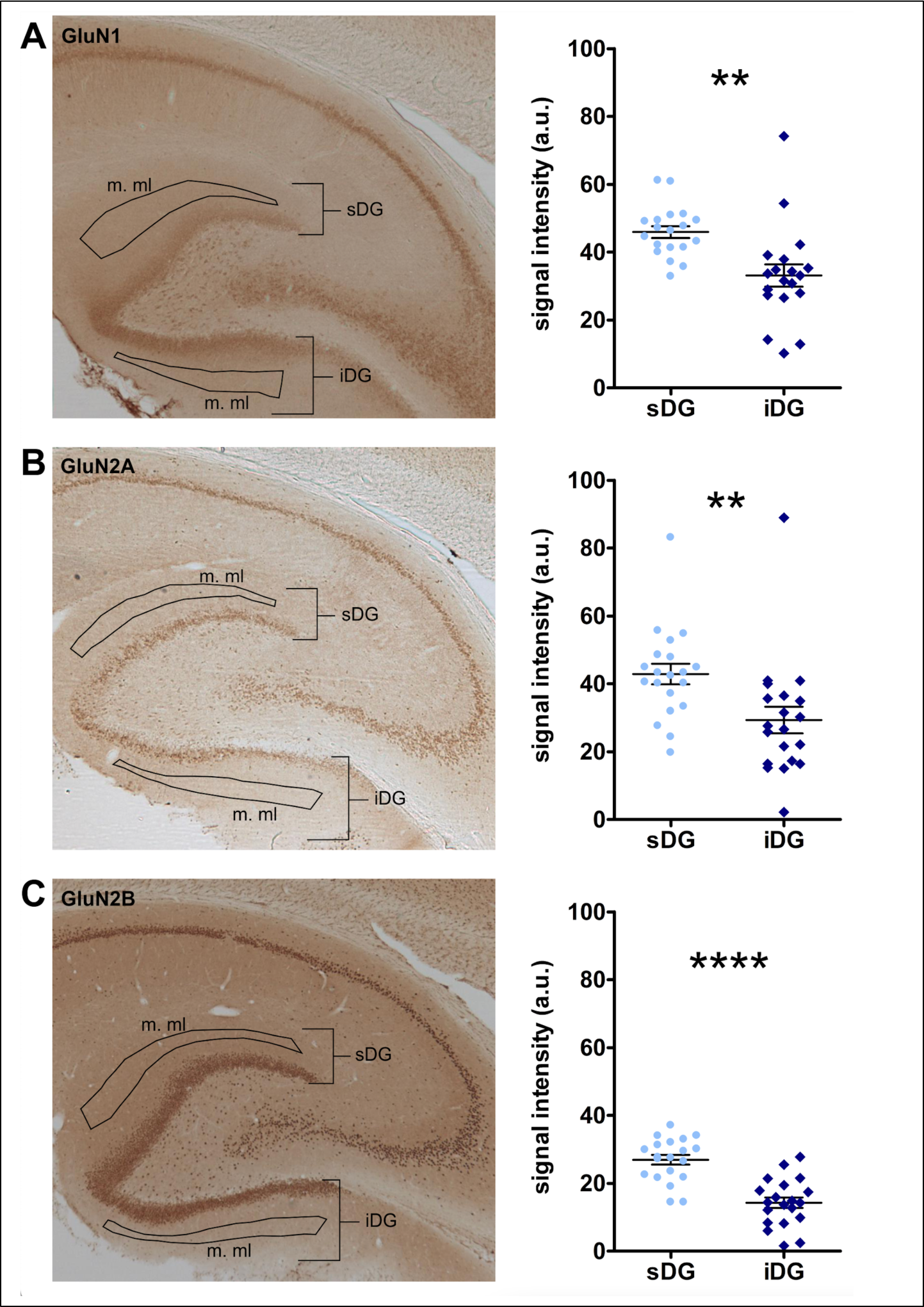
Expression of plasticity-related subunits of the NMDA receptor is lower in the middle molecular cell layer of the infrapyramidal compared to the suprapyramidal blade of the dentate gyrus. A) Examination of GluN1 subunits in the middle molecular layer (m. ml) reveals a significantly lower expression in infrapyramidal (iDG) compared to the suprapyramidal (sDG) blade of the dentate gyrus (N = 19 each). Example of a DAB-stained image of the hippocampal formation showing the expression of GluN1 subunit of the NMDA receptor. Outlines depict the regions of interest in the m. ml of the sDG and iDG. B) GluN2A expression is significantly weaker in the m. ml of the iDG compared to the sDG (N = 20 each). Expression of the GluN2A subunit in the hippocampal formation shown in an exemplary DAB-stained image. The regions of interest in the m. ml of the sDG and iDG are depicted within the outlines. C) In the m. ml of the iDG, GluN2B expression is significantly smaller than in the sDG (N = 20 each). Example of a DAB-stained photomicrograph showing the expression of the GluN2B subunit in a horizontal section of the hippocampus. The regions of interest in the m. ml are marked by outlines. A-C) Significant differences are marked with asterisks: ** p < 0.01; **** p < 0.0001.

## Discussion

This study is the first to compare frequency-dependent synaptic plasticity in MPP synapses of the iDG and sDG of the dorsal dentate gyrus of freely behaving rats. We observed that the magnitude of LTD induced by LFS (1 Hz), as well as of LTP induced by HFS (400 Hz) are weaker in the iDG compared to the sDG. HFS at 200 Hz induces no change in synaptic transmission in the iDG, whereas LTP in the sDG can be easily evoked by this frequency. Frequencies close to a-m (5 Hz and 10 Hz) (Kemp and Manahan-Vaughan, 2005; Twarkowski et al., 2016) applied to the MPP resulted in a transient depression followed by a late synaptic potentiation in both blades of the dorsal dentate gyrus. Patch clamp recordings revealed a smaller maximal firing frequency of granule cells of the iDG. Moreover, specific action potential properties differ between blades. Immunohistochemical analysis of the expression of NMDAR subunits in the middle molecular layer of the dentate gyrus where the MPP forms synapses (Steward, 1976; van Groen et al., 2002), revealed a lower amount of GluN1, GluN2A and GluN2B subunits in the iDG compared to the sDG. These results strongly support that both blades of the dorsal dentate gyrus exhibit unique properties, and different frequency-dependencies and expression profiles of synaptic plasticity that may form the basis for their functional differentiation with regard to spatial and non-spatial information processing (Chawla et al., 2005; Hoang et al., 2018; Vazdarjanova et al., 2006).

Until now, synaptic plasticity in the iDG of the granule cell layer of the dorsal dentate gyrus has not been examined in detail in behaving rodents. Many papers do not state which blade was targeted for recordings, but implantation coordinates, or histological evidence provided about electrode locations, indicate that most studies of synaptic plasticity in the dentate gyrus of behaving rodents focused on the dorsal sDG, whereas a few examined responses in the hilar region (Abraham et al., 1994; Altinbilek and Manahan-Vaughan, 2009; Blaise and Bronzino, 2003; Frey and Frey, 2009; Kenney and Manahan-Vaughan, 2013; Krug et al., 1983). In the present study, we specifically discriminated between the sDG and iDG and, as far as possible, conducted simultaneous recordings from both blades during MPP stimulation. We found that synaptic plasticity in the sDG induced by different afferent frequencies was comparable to previous studies, e.g. HFS at 200 Hz or 400 Hz applied to the MPP induce LTP in the sDG in behaving rats (Hansen and Manahan-Vaughan, 2015a; Klausnitzer et al., 2004; Manahan-Vaughan et al., 1998). By contrast, the iDG does not exhibit significant changes in synaptic strength following 200 Hz stimulation and although it expresses LTP in response to 400 Hz HFS the plasticity response is significantly smaller than LTP induced in sDG. In line with our findings, a study in anesthetized rats also reported the expression of LTP in the molecular layer of the iDG following 400 Hz stimulation of the angular bundle (Levy and Steward, 1979).

LTD was evoked in both blades by LFS at 1 Hz, although the magnitude of responses was weaker in iDG compared to sDG synapses. LFS at 5 or 10 Hz resulted in transient depression followed by slow potentiation in both sDG and iDG. These frequencies may lie within the range of a-m (Bienenstock et al., 1982; Twarkowski et al., 2016) that corresponds to afferent frequencies that are too high to induce LTD and too low to induce LTP. A clear result from this study is that long-term synaptic plasticity, in the form of LTD and LTP, are distinct in sDG and iDG, both in terms of their frequency-dependency and response profiles.

Scrutiny using patch clamp recordings revealed that the maximal firing frequency of granule cells in the iDG is indeed smaller compared to the sDG. Moreover, the threshold to induce an action potential is higher in the iDG, whereas the spike amplitude is smaller in iDG compared to the sDG. Other studies examined granule cell properties, but most of them focused on the depth of the respective cell within the granule cell layer, rather than a comparison of responses within the blades: For example, one study using whole cell recordings in vitro examined granule cells in the sDG and demonstrated that young cells deep in the granule cell layer (close to the hilar region) have a lower threshold for the induction of synaptic potentiation in contrast to more superficial cells (Wang et al., 2000). Another study using in vitro patch-clamp recordings compared three compartments (crest/hilus, sDG and iDG) of the dentate gyrus (Mishra and Narayanan, 2020). They did not identify distinct differences between iDG and sDG, but rather showed that properties within each region were heterogeneous (Mishra and Narayanan, 2020). Our results also reveal some heterogeneity across blades; but a comparison of results obtained in the two studies is difficult because of differences in the composition of aCSF and pipette solution. Interestingly, Mishra and Narayanan (2020) also examined the firing frequency of granule cells, but the maximal current they applied (250 pA) was in the range where in our study both blades still exhibited similar responses (Mishra and Narayanan, 2020) and the maximal frequencies of the currents < 250 pA they report are comparable to our own results. In our study differences first became apparent using higher currents (350-700 pA). It seems likely that the smaller maximal firing frequencies that we detected using higher currents as well as the higher threshold needed to induce an AP and the smaller AP spike amplitude in the iDG may create constraints with regard to the induction of synaptic plasticity and/or limit its magnitude (Stanton and Sejnowski, 1989; Wang et al., 1997). This may be particularly relevant for the induction of LTD in the dentate gyrus, especially considering that at least for the sDG it does not require activation of NMDAR (Poschel and Manahan-Vaughan, 2007). LTP responses are likely to have been strongly influenced by the weaker expression of GluN subunits of the NMDAR in iDG. In sDG, LTP induced by 200 Hz HFS is NMDAR-dependent in freely behaving rats, whereas LTP induced by 400 Hz stimulation recruits activation of both NMDAR and L-type voltage-gated calcium channels (VGCCs) (Manahan-Vaughan et al., 1998). Only 400 Hz HFS of the MPP resulted in LTP in iDG. LTP of different magnitudes and durations are induced by activation of GluN2A, or GluN2B, containing NMDAR (Ballesteros et al., 2016). Thus, the weaker LTP response of iDG may derive from the lower NMDAR content of this blade. One can only speculate as to whether L-type VGCCs contributed to LTP in iDG. Although we did not assess expression, others have reported that L-type VGCCs are needed for neurogenesis in the dentate gyrus (Marschallinger et al., 2015) and that the iDG exhibits higher neurogenesis than the sDG (Snyder et al., 2012), hinting that these channels must be present in iDG and thus could contributed to LTP in this blade.

In addition to the differences in cell properties and NMDAR receptor expression, other factors may contribute to the differences in synaptic plasticity between blades, such as distinctions in the morphology of the dendritic tree, inputs from the medial EC, or inhibitory control (Czéh et al., 2013; Woodson et al., 1989): For example, although it was shown that axons from the EC simultaneously branch and innervate DG granule cells of both blades (Tamamaki and Nojyo, 1993), other studies support that the medial EC projects preferentially to the iDG rather than the sDG, whereas the lateral EC projects preferentially to the sDG (Tamamaki, 1997; Wyss, 1981). This shift in input from the MPP to the different blades may play a role in the distinct profile of synaptic plasticity we detected. In addition, studies examining immediate early gene expression in the sDG and iDG revealed that exposure to a novel environment or novel spatial exploration of an environment results in an increase in Arc mRNA expression in the sDG, whereas no change can be detected in the iDG (Chawla et al., 2005; Vazdarjanova et al., 2006). In comparison, exploration of novel large environmental features results in an increase in Arc and Homer1a expression in the iDG, but not sDG (Hoang et al., 2018). These studies suggest that sDG and iDG play distinct and putatively separate roles in the processing of information obtained during novel exploration of environments and cues. This in turn may derive from differences in EC inputs to the blades.

The MPP input to the dentate gyrus forms synapses specifically on granule cell dendrites in the middle third of the molecular layer (Steward, 1976; van Groen et al., 2002), thus it is likely that a shift in the strength of the input from the MPP and/or LPP to the respective blades could also play a role in synaptic plasticity induced in both compartments of the granule cell layer. Morphological studies have shown that dendrites in the molecular layer of the sDG have greater length, more dendritic segments, a higher transverse spread and greater spine density compared to the molecular layer of the iDG (Claiborne et al., 1990; Desmond and Levy, 1985; Gallitano et al., 2016). One of these studies also examined dendrites in three compartments of the molecular layer (outer, middle and inner third) of the sDG and iDG in detail (Gallitano et al., 2016). They revealed that sDG granule cell dendrites in the middle molecular layer of rats contain a higher amount of dendritic material than iDG dendrites, albeit without exhibiting an increase in spine density or branching complexity (Gallitano et al., 2016). Thus, a general difference in dendrite morphology and spine density in the molecular layer, together with the higher amount of dendritic material in the middle molecular layer of the sDG may also impact on the ability of each blade to express distinct plasticity profiles.

Overall, the results of this study strongly support that the dentate gyrus can be subdivided into at least two functionally distinct cellular and synaptic compartments that differ in their basal excitability, NMDAR expression levels, and their ability to express long-term synaptic plasticity in the form of LTP and LTD. The differences in the profiles and frequency-dependency of persistent (>24h) forms of synaptic plasticity in sDG and iDG may reflect reported differences in information processing related to spatial and non-spatial experience. These findings also emphasize that the dentate gyrus cannot be considered as a uniform entity when deciphering its role in information processing and storage.

## Data availability statement

The data from this study are available from the corresponding author upon reasonable request.

## Acknowledgments

This project was supported by a grant from the German Research Foundation (Deutsche Forschungsgemeinschaft, www.dfg.de; SFB 1280/A04, project number: 316803389 to D.M.-V.). We gratefully thank Beate Krenzek and Ute Neubacher for technical assistance and Nadine Kollosch for animal care.

## Author contributions

The study was designed by D.M-V and further developed with C.S. In vivo electrophysiological experiments and analysis: J.B. and C.S., patch clamp recordings and analysis: O.S., immunohistochemistry and analysis: V.D. D.M-V. and C.S. interpreted the data and wrote the paper that was edited by all authors.

