## Supplemental Figure 1 for "The suprapyramidal and infrapyramidal blades of the dentate gyrus exhibit different GluN subunit content and dissimilar frequency-dependent synaptic plasticity in vivo"

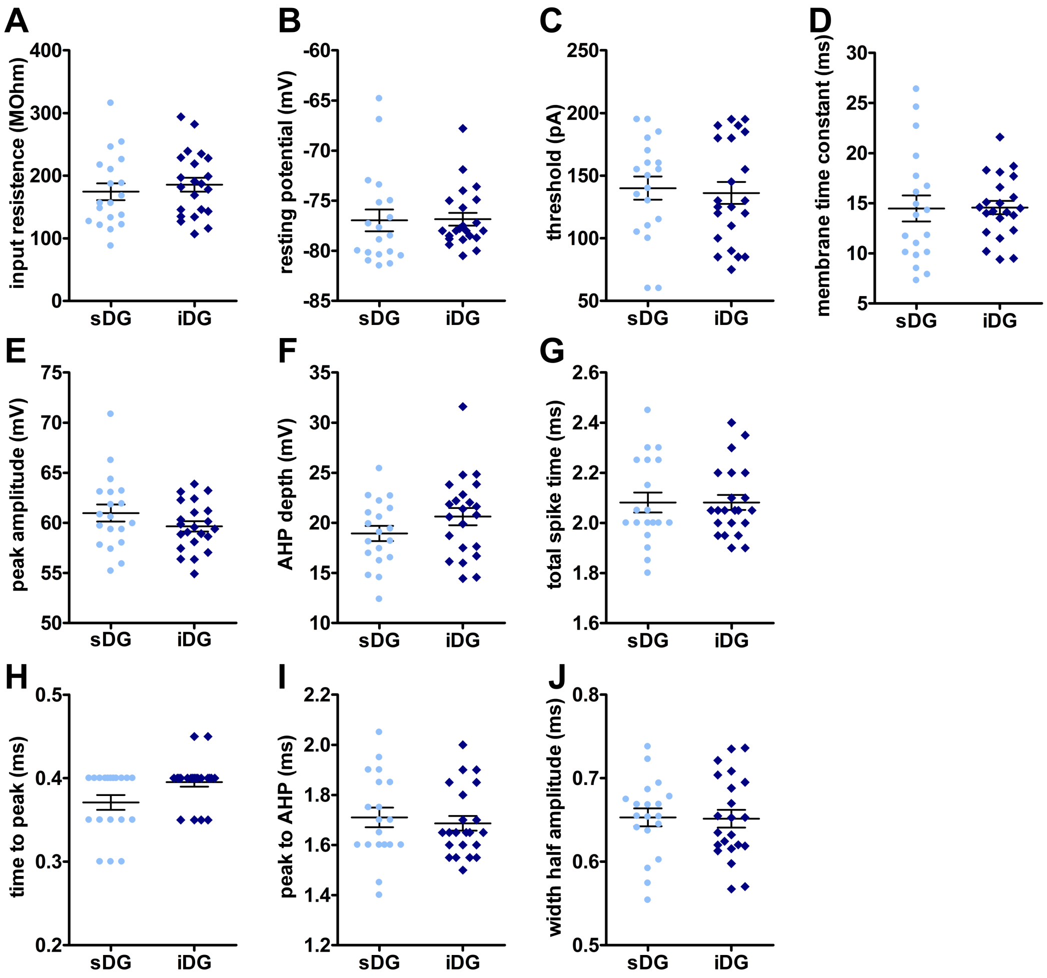


**Supplementary figure 1**

**Cell and action potential properties recorded in granule cells of the supra- and infrapyramidal blade during patch clamp recordings.**

A-D) Basic membrane properties of granules cells of the supra- (sDG) and infrapyramidal (iDG) blade of the dentate gyrus are not differing: A) The input resistance, B) the resting membrane potential, C) the threshold, and D) the membrane time constant (tau) are not different in granule cells of the sDG (n = 19, N = 6) and iDG (n = 22, N = 6).

E-K) Several action potential properties are similar in granule cells of the sDG (n = 19, N = 6) and iDG (n = 22, N = 6): E) The peak amplitude and F) the depth of the afterhyperpolarization (AHP) from the threshold are not different. G) The total spike time, H) the time from the threshold to the peak, I) the time from the peak of the action potential to the AHP, and J) the width at half of the maximum amplitude are comparable in granule cells of the sDG and iDG.
